# Super-dosed butyrate induces a revisable metabolic paralysis through transient mitochondrial reprogramming in the gut-brain axis

**DOI:** 10.1101/2020.12.23.424102

**Authors:** Yanhong Xu, Shiqiao Peng, Xinyu Cao, Shengnan Qian, Shuang Shen, Juntao Luo, Xiaoying Zhang, Hongbin Sun, Wei L. Shen, Weiping Jia, Jianping Ye

## Abstract

**Background and purpose:** Sodium butyrate (SB) is a major product of gut microbiota with signaling activity in the human body. However, the toxic effect of SB remains largely unknown. This issue is addressed in current study.

**Experimental approach:** SB (0.3 – 2.5 g/kg) was administrated through a single peritoneal injection in mice. The core body temperature and mitochondrial function in the brain hypothalamus were monitored. Pharmacodynamics, targeted metabolomics, electron microscope, oxygen consumption rate and gene knockdown were employed to dissect the mechanism for the toxic effect.

**Key results:** The temperature was reduced by SB (1.2 −2.5 g/kg) in a dose-dependent manner in mice for 2-4 hr. In the brain, the effect was associated with SB elevation and neurotransmitter (Glutamate and GABA) reduction. The mitochondria exhibited a transient volume expansion and crista loss in the hypothalamic neurons. ADP/ATP ratio was increased with accumulation of intermediate metabolites in the glycolysis, TCA cycle and pentose phosphate pathways. The mitochondrial protein, adenine nucleotide transporter (ANT), was activated for proton transportation leading to a transient potential collapse by proton leak. The SB activity was attenuated by ANT inhibition from gene knockdown or pharmacological blocker. The temperature drop was attenuated by i.p. injection of norepinephrine. The HDAC inhibitors, such as SAHA and pyruvate, did not exhibit the same effect.

**Conclusion and implications:** Super-dosed SB generated an immediate and reversible toxic effect for inhibition of body temperature through transient mitochondrial reprogramming in the brain. The mechanism was quick activation of ANT proteins for the proton leak in mitochondria.

## 1. Introduction

Sodium butyrate (SB) is a signaling molecule in the gut-brain axis (De Vadder et al., 2014), by which the gut microbiota generates an impact in the brain in the regulation of energy metabolism (Bienenstock, Kunze & Forsythe, 2015; Canfora, Jocken & Blaak, 2015). SB is a natural product of bacterial fermentation of dietary fibers in the large intestine, which mediates the signal of gut microbiota to keep energy balance in the host body in favor of control of obesity, type 2 diabetes, hepatic steatosis, etc. (Canfora, Jocken & Blaak, 2015; Forslund et al., 2015; Qin et al., 2012). The mechanisms are related to induction of energy expenditure by activation of PGC-1α and AMPK for mitochondrial biogenesis (Gao et al., 2009), by induction of FGF21 for fat utilization (Li et al., 2012) and inhibition of food intake for calorie restriction (Li et al., 2017). In addition, SB plays a role in the maintenance of intestine function, training of T cells in control of autoimmune reactions (Donohoe et al., 2011; Furusawa et al., 2013). In the brain, SB protects neurons from ischemia damage, improve long-term memory, attenuate neurodegenerations and even inhibits brain tumor growth (Stilling, van de Wouw, Clarke, Stanton, Dinan & Cryan, 2016). As a histone deacetylase inhibitor (HDACi), SB has an activity in the treatment of brain diseases, such as Huntington’s Disease and Alzheimer’s disease (Stilling, van de Wouw, Clarke, Stanton, Dinan & Cryan, 2016). Derivatives of SB (such as β-hydroxybutyrate) have fruit flavors, which are widely used as additives in the sport drinks. Some forms of butyrate are dietary supplements for obesity subjects (Roberts et al., 2014) and for patients with functional disorders of large intestine (Canfora, Jocken & Blaak, 2015; Rojas-Morales P, Tapia E & J., 2016). With the broad applications, there is a clear demand for understanding of the over-dosed effects of SB. Unfortunately, there is little information about the SB effect over 1.2 g/kg (body weight) in the literature (Stilling, van de Wouw, Clarke, Stanton, Dinan & Cryan, 2016). This study was conducted to test the immediate effect of SB at dosages above 1.2 g/kg.

SB is actively taken up in the brain and metabolized in the neuronal cells (Li et al., 2019; Li et al., 2017; Val-Laillet et al., 2018). SB enters the mitochondria through the protein channel of MCT1 and converts into acetyl-CoA in the TCA cycle for ATP production, which fuels cells in the brain and other tissues (Kim et al., 2013; Li et al., 2019). In addition to the energy supply, SB has a well-known activity of HDACi in the control of transcription of nucleus-encoded genes, an epigenetic effect. The genes, such as PGC-1α, is known to promote mitochondrial biogenesis (Gao et al., 2009). These activities of SB have been documented in the studies of nutritional and pharmacological activities of SB (Mollica et al., 2017; Walsh et al., 2015). However, the immediate impact of overdosed SB in mitochondria remains unknown in the gut-brain axis.

ANT (ADP/ATP carrier, AAC in yeast) is a nucleus-encoded protein located in the inner membrane of mitochondria. The traditional activity of ANT is import of ADP and export of ATP in mitochondria in the control of ATP production and supply. ANT is a component in the mitochondrial permeability transition pore complex originally identified in the study of cell apoptosis (Zamzami et al., 1996). The pore complex contains three isoforms of ANTs (ATN1, ANT2 and ANT4) (Karch et al., 2019). ANT has recently been identified as a new class of proton transporter, whose activity is induced by the long chain fatty acids in the brown adipocytes (Bertholet et al., 2019). Pharmacological inhibition of ANT activity promotes mitophagy independently of the traditional activity in ADP/ATP transportation (Hoshino et al., 2019). Over activation of ANT leads to a proton leakage for the collapse of mitochondrial potential in the mechanism of cell apoptosis (Qin et al., 2020). This impact on mitochondria may mediate the SB effects in the brain, especially at the high dosages. However, the possibility has not been tested by experiment.

We investigated the immediate impact of super dose SB in the brain. SB induced a reversible inhibition of mitochondrial function in the brain, which led to a reduction in the body energy metabolism with a fall in the body temperature. The temperature change was reversed partially be the injection of norepinephrine.

## 2. Materials and Methods

### 2.1 Mice and SB treatment

Male C57BL/6 mice at 6-8 weeks of age were purchased from the animal facility of Nanjing University, and maintained in the animal facility (SPF) of the Shanghai Sixth People’s Hospital, Shanghai Jiao Tong University, with 12 hr light cycle, temperature of 22 ± 2 °C, humidity of 60 ± 5%, free access to water and Chow diet (13.5% calorie in fat; Shanghai Slac Laboratory Animal Co). All animal procedures were conducted according to the animal protocol approved by the Institutional Animal Care and Use Committees (IACUC) at the Shanghai Sixth People’s Hospital, Shanghai Jiao Tong University. In the SB treatment, the mice were fasted for 10 hr and then injected i.p with 200 μl SB solution (S817488, MACKLIN,China) at dosages as indicated in the figure legend. The control mice were injected with the same volume of saline. In the comparison study, sodium chloride, sodium pyruvate, and sodium succinate were used at the same dosage in molar.

### 2.2 Core body temperature and pharmacodynamics assay

The core body temperature was measured with an anal thermometer (Type C Thermometer, Therma Waterproof) under conditions as indicated. The skin temperature was measured using the thermal infrared imager (Smart Sensor, China). Blood, brain tissues and brown adipose tissue (BAT) were collected immediately after the cervical dislocation at different time points (10 - 240 min) after the SB injection (2.5 g/kg body weight) and kept at −80 °C until the measurement. SB was extracted from the samples and measured with GC-MS (Agilent 7890B). NaCl, sodium pyruvate, beta-hydroxybutyrate, and sodium succinate were used in the controls for sodium or fatty acid salts. SAHA was used at 50 mg/kg as a control of histone deacetylase inhibitor.

### 2.3 Transmission electron microscopy

The fresh brain tissue was collected at different time points (0 – 240 min) as indicated in the figure legend after SB (2.5 g/kg) injection. The hypothalamic tissue (1 mm^3^) was fixed in 2.5% glutaraldehyde solution at room temperature for 2 hr and then 4 °C for 4 hr. The specimen was fixed in 1 % osmium acid, dehydrated with gradient ethanol, embedded in epoxy resin and sectioned in ultrathin (60 – 80 nm, Leica, UC7). The mitochondrial ultrastructure picture was taken with a transmission electron microscopy (HITACHI, HT7700).

### 2.4 Targeted metabolomics

The fresh brain tissue was collected at 30 min after SB injection (2.5 g/kg, i.p.) and frozen in liquid nitrogen immediately and kept at −80 °C until analysis. The metabolites (30 small molecule chemicals) in the glycolysis, TCA cycle, oxidative phosphorylation, and pentose phosphate pathways were analyzed using the ultra-performance liquid chromatography (Agilent 1290 Infinity LC) coupled with triple quadrupole mass spectrometry (5500 QTRAP, AB SCIEX). The hemisphere tissue (about 100 mg) was homogenized in ultra-pure water, treated with mixture of methanol and acetonitrile (1:1 v/v), and centrifuged at 14000 g at 4 □ to remove protein after incubation at −20 □ for 1 hour. The supernatant was dried in vacuum and used for detection of metabolites in a mixture solution of acetonitrile and water (1:1, v/v). The data was analyzed with Multiquant program against the standard compounds.

### 2.5 Neurotransmitter assay

The samples collected at 30 min after SB (2.5 g/kg) injection was homogenized in the analysis of glutamate and γ-aminobutyrate (GABA). The homogenization was prepared in a mixture of methanol, acetonitrile and water (2:2:1, v/v/v) and then diluted with the same volume of chloroform. The supernatant was collected for detection of glutamate and GABA. Analyses were carried out using ultra-performance liquid chromatography (H-Class UPLC) coupled with Xevo TQ-XS triple quadrupole MS (Waters, Milford, MA, USA). In the plasma assay, the blood was collected from eye vein with EDTA anticoagulant tube, and the neurotransmitters were quantified in the plasma with LC-MS. The small molecules were separated with Agilent 1290 Infinity LC and their mass were examined with 5500 QTRAP mass (AB SCIEX). The data was analyzed with the MultiQuant software program against the standards.

### 2.6 OCR assay of isolated mitochondria

Mitochondria were extracted from the brain hemisphere homogenization in the MSE buffer (210 mM mannitol, 70 mM sucrose, 1 mM EGTA and 0.5% BSA) using a protocol in our early study (Sun, Jin, Zhang, Jia, Le & Ye, 2018). The oxygen consumption rate (OCR) was detected in the mitochondria (10 ug/well) with the Seahorse XFe24 (Seahorse Bioscience, Agilent, Santa Clara, USA) to detect the coupling capacity. SB (10 mM or 20 mM) was administrated to the mitochondria before the test. The substrates in MAS were 5 mM pyruvate and 10 mM succinate. The activators and inhibitors were as following: ADP (4 mM, A5285, Sigma), Oligomycin (Oligo, 1 μM, 495455, Sigma), FCCP (2 μM, C2920, Sigma), and Antimycin A (AA, 0.5 μM, 2247-10, Biovision).

### 2.7 ANT knockdown

To reduce ANT1 and ANT2 in the cell line of Neuro-2a, we designed a shRNA targeted to ANT1 with adenovirus as vector (pADV-U6-shRNA (Slc25a4)-CMV-EGFP). The target sequence was: GCAGTTCTGGCGCTACTTT, which has a high degree of homology with ANT2 for one base pair difference. Both ANT1 and ANT2 were effectively knocked down in the Neuro-2a by this shRNA. The stock solution of virus was 5.53 × 10^10^ PFU (plaque forming unit)/ml in titer. The MOI (Multiplicity of infection) was 100 IFU (infect formation unit)/cell in the experiment.

### 2.8 WB, qRT-PCR and cell culture study

Supplement 1.

### 2.9 Statistical analysis

The in vivo data were statistically analyzed with the two-way ANOVA and the in vitro data were analyzed with Student T-test. In vitro, all experiments were repeated at least three times with consistent results. The data are presented as mean ± SEM with a significance of *p*<0.05.

## 3. Result

### 3.1 SB reduces body temperature in a dose-dependent manner

SB has been used as a HDAC inhibitor in various disease models including cancer, in which the highest dosage is 1.2 g/kg (Stilling, van de Wouw, Clarke, Stanton, Dinan & Cryan, 2016). The effect of SB above the dosage remains largely unknown, especially in the regulation of energy metabolism. To address the issues, SB was administrated in the healthy mice through a bolus i.p. injection at dosages 0.3 – 2.5 g/kg in this study. The core body temperature and behavior were monitored in those mice at the ambient room temperature. SB induced a transient and reversible drop in the body temperature in a dose-dependent manner at dosages above 1.2 g/kg including 1.5 g/kg, 1.8 g/kg and 2.5 g/kg (Fig. 1A). The peak drop was observed at 1 hr post injection with 6 °C below the normal body temperature, which was associated with a skin temperature drop (Fig. 1, A and B). The temperature recovered to the normal level after 4 hr post injection. At the dosage of 5 g/kg, the temperature drop became irreversible leading to mortality within 6 hr post injection (Fig. 1C). The speed of temperature drop was affected at the ambient temperature. The temperature drop was accelerated at 4 °C but attenuated at the neutral environment temperature of 29 °C (Fig. 1D). To test the impact of sodium ion in the molecule of SB, sodium chloride (NaCl) and sodium pyruvate (SP) were used at the same dosage for a comparable load of sodium ion in the mice. The temperature was modestly increased by sodium chloride and modestly decreased by sodium pyruvate around 1 °C (Fig. 1E). β-hydroxybutyrate (BHB) and succinate (SUCC) were compared with SB for the similarity in carbon bone length. β-hydroxybutyrate induced a small drop of 2 °C and succinate reduced the temperature with a similar potency to SB (Fig. 1E). In the behavior, the mice exhibited much less locomotion and showed a slow response to pinch challenge, which disappeared after the temperature recovery. These data suggest that the temperature drop is one of the immediate impacts of high dose SB above 1.2 g/kg in the bolus injection together with a reduction in movement and responses. The thermogenic function was inhibited by SB as indicated by the impact of ambient temperature. The suppression is related to the fatty acid structure, but not to the sodium ion in the SB molecule.

**Fig. 1.**
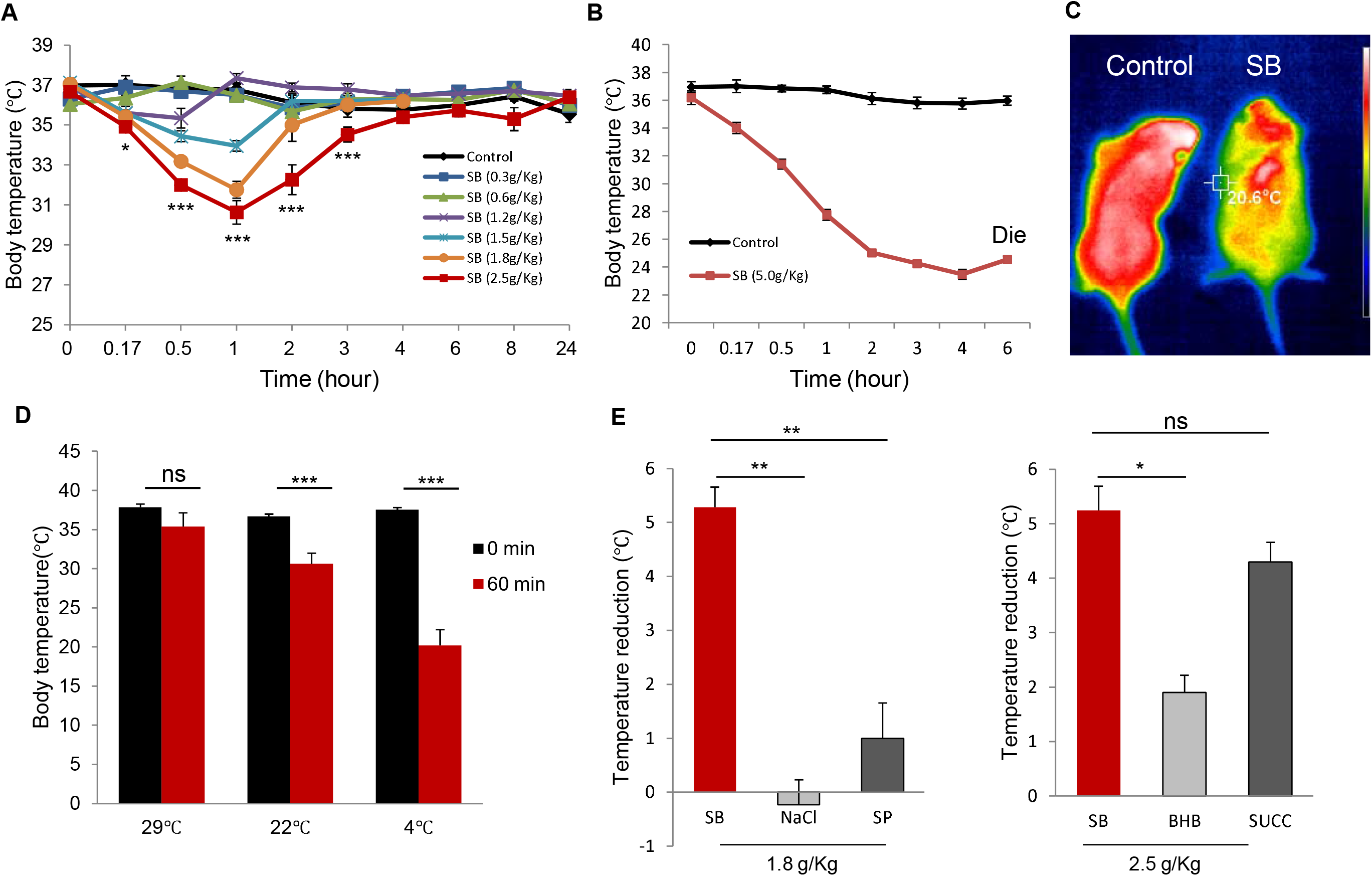
Reduction of body temperature by SB in a dose-dependent manner. (A) Sodium butyrate (SB) reduced the body temperature at dosages of 1.2 – 2.5 g/kg by intraperitoneal injection (i.p.) (n=6). (B) Toxicity of SB at 5.0 g/kg. This dosage reduced the body temperature more effectively and killed all of the mice at 6 hr (n=6). (C) Infrared imaging of body temperature. The image was taken at 30 min of SB (2.5 g/kg) injection. (D) Body temperature at different ambient temperature (n=3). (E) Substance comparison. Different substances were applied to the mice i.p. at the same dosage in molar. and the temperature was determined at 1 hour of injection. In comparison of NaCl and SP, the dose was 1.8 g/kg. In comparison of BHB and SUCC, the dose was 2.5 g/kg. The data value represents mean ± SD. SB (n=6), NaCl (sodium chloride, n=3), SP (sodium pyruvate, n=3), BHB (β-hydroxybutyrate, n=3), SUCC (sodium succinate, n=4). * *p*<0.05, ** *p*<0.01, *** *p*<0.001 vs control.

### 3.2 Pharmacokinetics of SB in the brain and blood

Above data suggest that SB may generate the effects through an impact in the brain, which is in line with the uptakes and metabolism of SB in the brain (Li et al., 2019; Li et al., 2017; Val-Laillet et al., 2018). However, the pharmacokinetics of SB remains unknown in the brain. To address this issue, SB concentration was examined in the brain and blood following the SB administration. SB increased quickly in the blood with a peak of 12 mM at 10 min and decreased quickly thereafter (Fig. 2A). The half-life was 60 min in the blood. A similar pattern of pharmacokinetics was observed in the brain while the peak duration was longer (Fig. 2B), which was established at 10 min and lasted for 20 min. The half-life of SB was 120 min in the brain. In the peripheral tissue, SB was determined in the brown fat tissue. The peak was at 30 min and half-life for 90 min (Fig. 2C). The elevation became not detectable at 240 min (4 hr) in all tissues. The brain pharmacokinetics matched the dynamics of temperature fall and recovery in the mice.

**Fig. 2.**
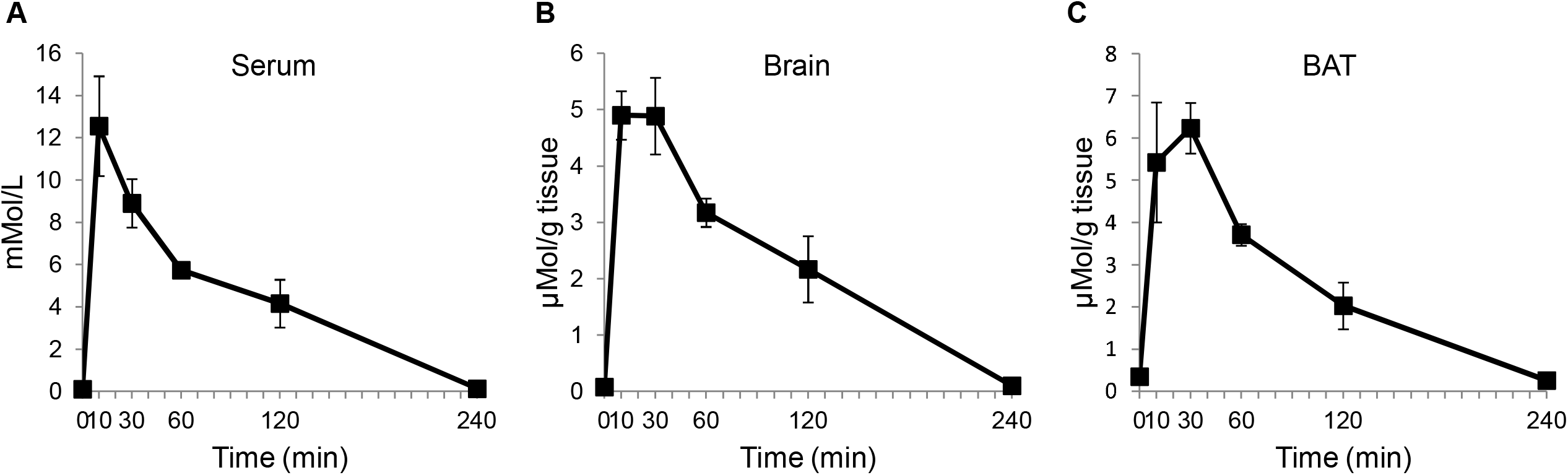
Pharmacokinetics of sodium butyrate after SB intraperitoneal injection. Dose of SB is 2.5 g/kg (n=3) at each time point. The SB concentration was detected with the method of GS/MS.

### 3.3 SB reduces neurotransmitters in the brain

Neurotransmitters mediate the central nervous signals in the control of thermogenesis of peripheral tissues. Above data suggests a brain inhibition by SB. To test the possibility, the representative neurotransmitters, GABA and glutamate, were examined in the brain tissue at 30 min of SB injection. Their concentrations were significantly reduced in the brain (Fig. 3A), as well as in the blood (Fig. 3B). Other neurotransmitters including histamine and glutamine were also reduced in the blood. However, the hormone epinephrine was elevated by two folds. No change was found in serotonin, tyramine, and 5-HIAA. The sympathetic nerves release norepinephrine to activate the brown fat in the cold response. Norepinephrine was not detected in the plasma, but its derivative, normetanepherine was reduced in the plasma. The cAMP/PKA signaling pathway was down regulated at the downstream of norepinephrine β3 receptors in the brown fat for a reduction in pCREB and pHSL (Fig. 3, C and D). Norepinephrine injection partially reverse the temperature drop and OCR inhibition by SB in vivo (Fig.3, E and F), which was observed with an increase in pHSL in the brown fat mitochondria in the SB-treated mice (Suppl. 2B). These data suggest that both the central and peripheral nervous system are inhibited by SB for the temperature drop.

**Fig. 3.**
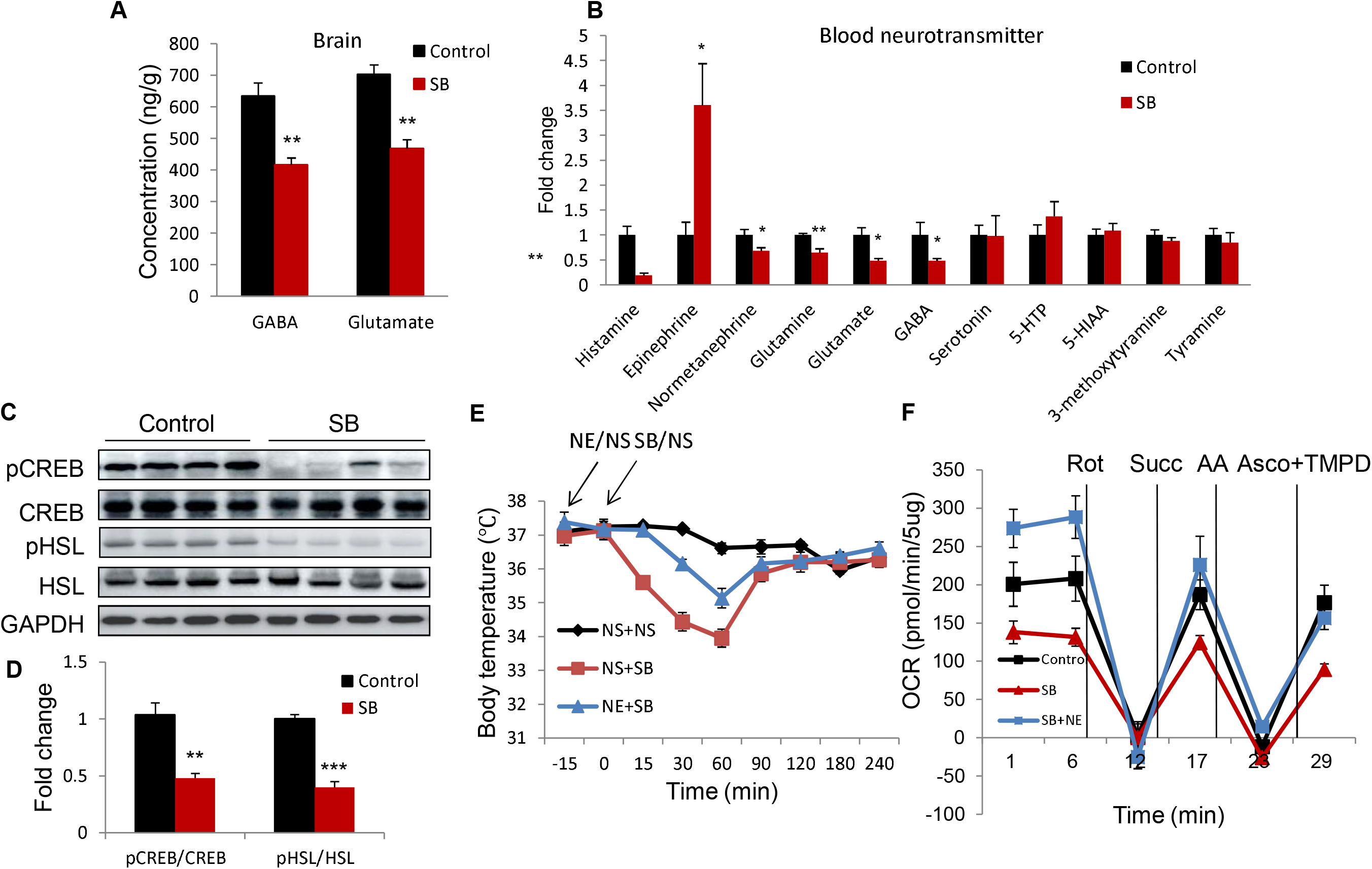
Changes of neurotransmitters in the brain and plasma. (A) GABA and glutamate were reduced in brain after 30 min of SB injection (2.5 g/kg, n=6). GABA: gamma amino butyric acid. GABA and glutamate were detected with method of LS/MS. (B) The change of neurotransmitters in plasma after 30min of SB i.p (2.5 g/kg, n=6). Neurotransmitters were detected with method of LS/MS. (C) Western blot showed the change of phosphorylation state of HSL and CREB in brown fat after 1 hour of SB injection (n=4). (D) Signal quantification. The blot signal in the panel C was quantified and presented after normalization with protein loading. (E) Norepinephrine (NE) effect. NE partially reversed the temperature drop in the SB model (n=7). (F) NE reversed the reduction of OCR caused by SB. The dose of SB was 2.5 g/kg. NE was 1 mg/kg administrated by i.p. at 15 min before SB. Mice were sacrificed at half hour after SB injection and brown fat mitochondria were used in the test (n=3). The data value represents mean ± SD. \**p*<0.05, \*\**p*<0.01, \*\*\**p*<0.001 vs control.

### 3.4 SB induces a transient mitochondrial stress in the brain hypothalamus neuron

SB is metabolized in mitochondria through several steps including β-oxidation, TCA cycle and oxidative phosphorylation in production of ATP. Mitochondrial failure is a common risk factor for the neuronal dysfunctions in conditions including poisoning, ischemia, and hypoxia. Mitochondrial super structure was examined in the brain hypothalamus neuron under the electron microscopy to understand the function. In the SB-treated mice, the mitochondria exhibited swelling with an increase in size and loss of crista number in the SB group (Fig. 4A). The changes lasted for 30 - 60 min and disappeared at 240 min post the SB injection, which was consistent with the temperature recovery in the SB group.

**Fig. 4.**
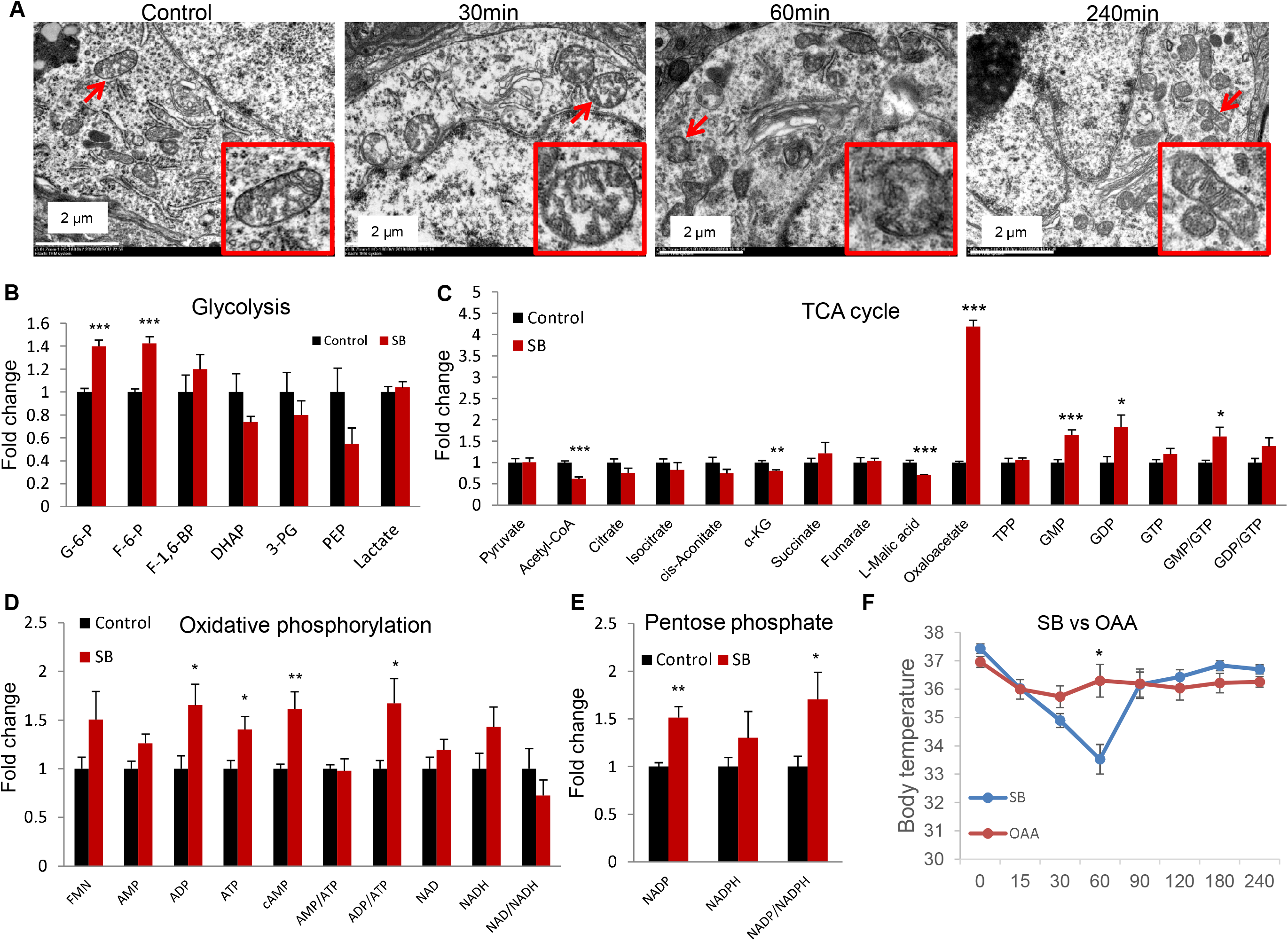
Mitochondrial structure and metabolism changes in the brain after SB injection. (A) Mitochondria swelling and crista breakdown were found at 30 −60 min in the hypothalamus neuron after SB injection at the dose of 2.5 g/kg. (B)-(E) Metabolite profile change in the brain at 30 min after SB (2.5 g/kg) administration (n=6). The changes of metabolites were found in the pathways of glycolysis, TCA cycle, oxidative phosphorylation and pentose phosphate. (F) Effect of oxaloacetate (OAA) in the regulation of body temperature. OAA was used at the dosage of 1.8 g/kg identical to that of SB in this study (n=6). The value represents mean ± SD. \**p*<0.05, \*\**p*<0.01, \*\*\**p*<0.001 vs control.

Metabolomics and oxygen consumption rate (OCR) were examined in the functional study of mitochondria. The energy-related metabolites were determined in the brain tissue by the targeted metabolomics. An increase was observed in the glycolytic intermediates, such as G-6-P and F-6-P (Fig. 4B). A mixed changed was observed in the TCA cycle metabolites with an elevation in oxaloacetate, a decrease in acetyl-CoA, malic acid and α-ketoglutarate (Fig. 4C). In the parameters of energy status, ATP and ADP were increased together with the ADP/ATP ratio (Fig. 4D). Similarly, an elevation was observed in GDP and GMP with the increases in the ratios of GMP/GTP, and GDP/GTP (Fig. 4D). These happened with an elevation in the brain cAMP level (Fig. 4D). In the pentose phosphate pathway, an elevation in NADP was observed leading to an increase in the ratio of NADP/NADPH (Fig. 4E). The 4-fold increase in oxaloacetate was the most impressive response in the metabolites. The role of oxaloacetate was unknown in the SB activity. To address the issue, oxaloacetate was administrated in the mice and a minor temperature drop was observed (Fig. 4F), which was much weaker than that of SB at the same dosage. These data suggest that the mitochondrial function was inhibited by SB in the brain in a reversible manner. Oxaloacetate elevation may mediate a part of SB effect. The oxaloacetate elevation may be a consequence of feedback inhibition by the elevated ATP.

### 3.5 SB reduces mitochondrial potential through MPTP-mediated proton leak

The structural damage and metabolic alterations of mitochondria suggest a loss of mitochondrial potential (Δψm). To test the possibility, Δψm was examined in the isolated mitochondria of brain tissue. A significant reduction was observed in the SB-treated mice (Fig. 5A). Δψm reflects the balance of respiration and proton leak. The mitochondrial respiration was increased by SB for the elevated OCR (Fig. 5B), which was blocked by the ATPase inhibitor (Oligomycin). The proton leak was induced by SB (Fig. 5B). These data suggest a role of proton leak, not respiration inhibition, in the Δψm reduction by SB.

**Fig. 5.**
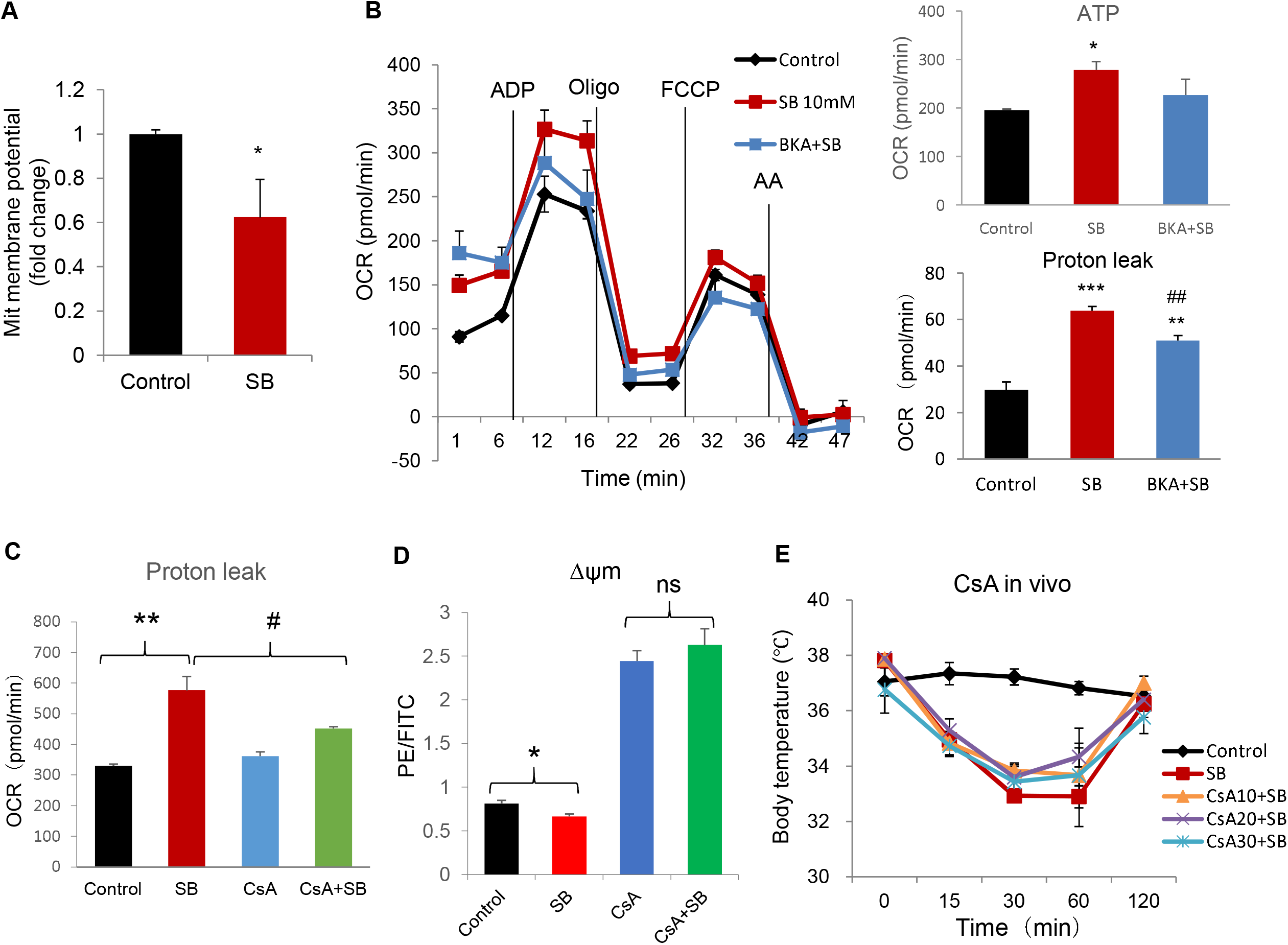
SB reduces brain mitochondrial membrane potential by activation of ANT. (A) SB reduced brain mitochondrial membrane potential. Brain mitochondria were extracted 1 hour after SB (2.5 g/kg) injection. JC-1 was used to detect mitochondrial membrane potential with the concentration of 2 µg/ml (n=8). (B) ANT inhibitor bongkrekic acid (BKA) reduce the proton leak induced by SB in vitro. Mouse hemisphere mitochondria were extracted and then treated with SB in vitro before OCR assay using the Seahorse equipment. The SB dose was 10 mM. The dose of BKA was 5 µM. The substrates were MAS with 10 mM succinate and 2 µM rotenone with 10 µg mitochondria per well (n=3-4). (C) Cyclosporin A ameliorated SB effect in the induction of proton leak in neuro2A cells detected by seahorse. CsA was 5 μ M,and SB was 20 mM in this experiment (n=3-4). (D) CsA blocked potential drop induced by SB. In the neuro2A cells test, SB was 20 mM and CsA was 5 μM. The potential was determined with JC-1 by flowcytometry (n=3). (E) Cyclosporin A (CsA) failed to reverse the reduced body temperature in vivo. CsA was injected daily for 3 days at three doses (10 mg/kg, 20 mg/kg, 30 mg/kg i.p.). SB was injected at the dose of 1.5 g/kg after 1 hour of the third injection of CsA (n=3-4). There were no differences in groups with or without CsA. Comparing with control, * p<0.05, ** p<0.01, *** p<0.001 vs control, ## p<0.01 vs SB.

Opening of the mitochondrial permeability transition pore (MPTP) in the inner mitochondrial membrane (IMM) is a mechanism of proton leak. ANT1 and ANT2 (AAC, ADP/ATP carrier in yeast) are proton transporters in the pore complex (Bertholet et al., 2019; Zhivotovsky, Galluzzi, Kepp & Kroemer, 2009). The ANT activity was blocked with the specific inhibitor (BKA), which significantly reduced the proton leak induced by SB (Fig. 5B). To confirm the result, Cyclosporin A (CsA), a MPTP inhibitor (Zhivotovsky, Galluzzi, Kepp & Kroemer, 2009), was used and the proton leak was significantly reduced (Fig. 5C, and Suppl. 3). The potential drop induced by SB was also blocked by CsA (Fig. 5D). However, in vivo, CsA injection failed to block the temperature drop in the SB-treated mice (Fig. 5E). Peripheral CsA is not able to cross the blood-brain barrier (Matsumoto, Murozono, Kanazawa, Nara, Ozawa & Watanabe, 2018). The data suggest that SB may activate ANTs to induce the transient mitochondrial potential collapse in the brain.

### 3.6 ANT mediates the SB activity in mitochondria

The MPTP complex is formed by multiple proteins, such as ANT1, ANT2, VDAC, TOMM20, Bcl-2, Bcl-xL, Bax and Bad (Karch et al., 2019; Zamzami et al., 1996). These proteins were not significantly changed in the SB-treated brain except a weak reduction in ANT1 (Fig. 6A). There are two isoforms of ANTs (ANT1 and ANT2) in the neuronal cells (Karch et al., 2019). ANT1 was three folds of ANT2 in the expression level (Fig. 6B). The ANT activity was further investigated with shRNA-mediated knockdown (Fig. 6 C). The knockdown did not inhibit the ATP production or proton leak in the absence of SB in the neuronal cells of Neuro2a (Fig. 6D). However, it blocked the SB-induced ATP production and proton leak (Fig. 6D). Both ATP production and proton leak were increased by SB before the knockdown. These results suggest that ANTs mediate the SB activity in proton leak.

**Fig. 6.**
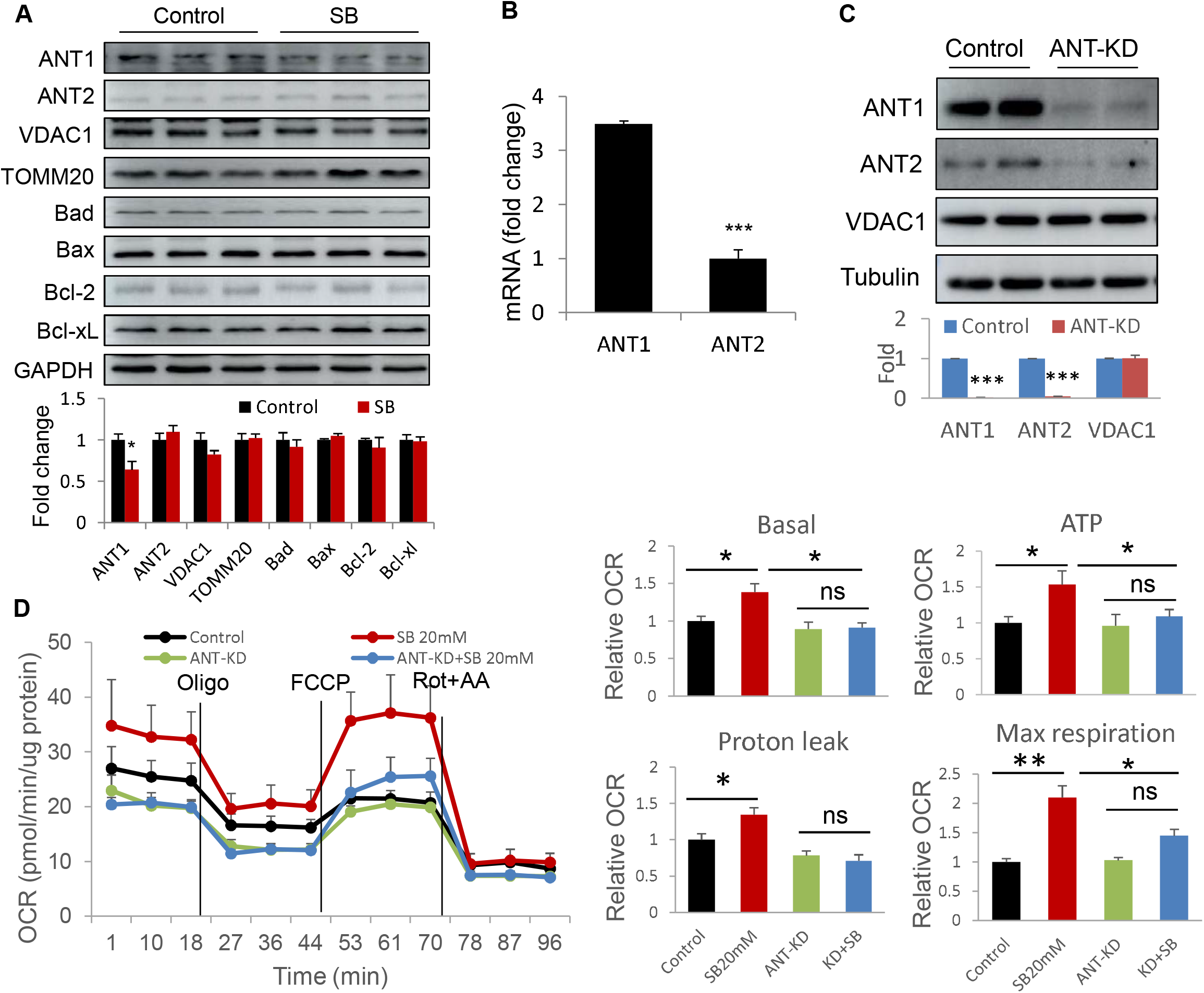
SB effect is dependent on ANT proteins. (A) Representative protein in mPTP complex. WB was conducted in the brain tissue of SB-treated mice. (B) ANT1 and ATN2 in the mouse brain tissue. ANT1 was about 3.5 folds of ANT2 in mRNA level determined with qRT-PCR (n=6); (C) Knockdown effect. shRNA-mediated knockdown of ANT1 and ANT2 in the brain cells with protein reduction by 90%; (D) The SB effect was abolished by ANT knockdown (n=3). In this study, each experiment was repeated three times with consistent results. The representative blot and OCR curve are shown. In the Western blot, each lane represents one mouse. * *p*<0.05, *** *p*<0.001 vs control.

### 3.7 Gene expression may not play a major role in the SB activity

Gene expression was investigated to understand the HDACi activity of SB. The histone acetylation was induced by SB time-dependently after SB injection (Fig. 7A). Global acetylation was increased as well in the brain tissues by SB (Fig. 7, B and C). Expression of representative genes was examined for the TCA cycle, mPTP complexes and receptors of short chain fatty acids. mRNA expression of some of them (CS, OGDH, SDH, ANT1, ANT2, GPR41 and GPR43) was induced by SB at 1 hr (Fig. 7D). However, the change did not lead to protein alteration in the TCA cycle and respiratory complexes at the same time (Fig. 7, E and F). The representative HDACi, SAHA, was used in the control and the effect on the body temperature was much weaker that of SB (Fig. 7G). These data suggest that SB modulated mRNA expression through HDACi. However, the gene expression may not play a major role in the SB activity in the brain.

**Fig. 7.**
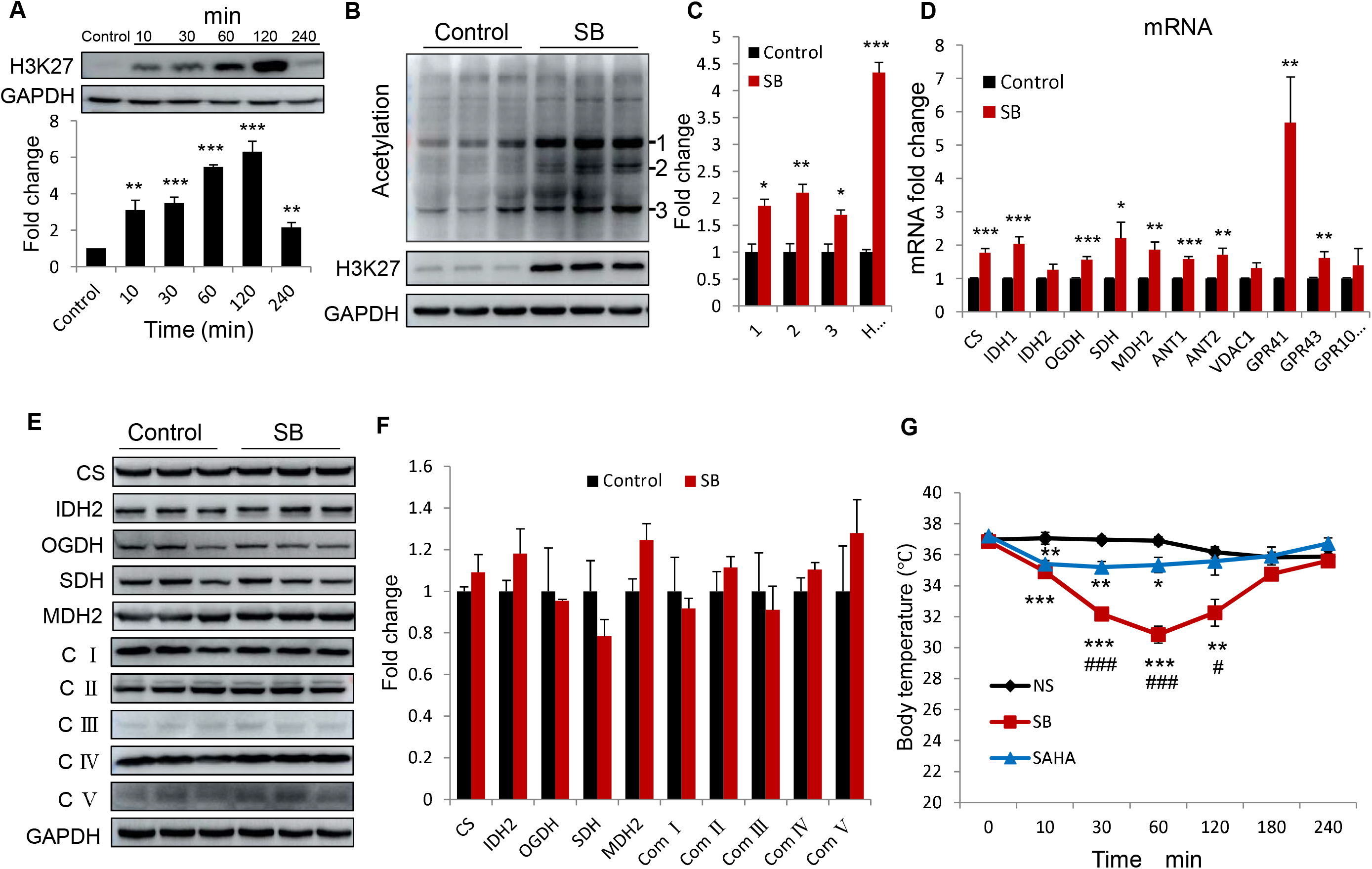
SB increases protein acetylation without alteration of protein abundance of mitochondrial enzymes. (A) SB increases H3K27 acetylation of histone protein in the brain tissue in a time-dependent manner, with the highest acetylation state at 2 hours of SB injection (2.5 g/kg) (n=3). (B) SB increased global acetylation in the brain. The tissue was collected at 1 hr of SB administration (n=3). (C) Signal quantification of the panel B. The global acetylation signal was normalized with protein loading. (D) Gene expression determined by qRT-PCR. The test was performed at 1 hr of SB treatment (n=6-8). CS: Citrate synthase, IDH1: Isocitrate dehydrogenase 1, OGDH: Ketoglutarate. dehydrogenase, SDH: Succinate dehydrogenase, MDH2: Malate dehydrogenase 2. (E) and (F) Protein in the brain after 1 hr of SB administration. (G) HDACi. The representative HDACi, SAHA, decreased the body temperature at the dose of 50 mg/kg, but the effect was significantly weaker than that of SB (n=6). The in vitro experiments were repeated three times with consistent results. Representative blots are shown. * p<0.05, ** p<0.01, *** p<0.001 vs control. # p<0.05, ## p<0.01, ### p<0.001 vs SAHA.

## 4. Discussion

This study provides a new insight into the mechanism of SB action in the gut-brain axis. SB was tested at super dosages (1.5-2.5 g/kg) relative to the widely-used dosage of 1.2 g/kg in the pharmacological studies (Stilling, van de Wouw, Clarke, Stanton, Dinan & Cryan, 2016). In the rodent models, SB benefits neuronal plasticity, long-term memory, treatment of neurodegeneration diseases, and protection from the traumatic brain injury (Stilling, van de Wouw, Clarke, Stanton, Dinan & Cryan, 2016). SB induces a stress like-response in the hypothalamus and anxiety-like behavior by i.p. injection at 1.2 g/kg (Gagliano, Delgado-Morales, Sanz-Garcia & Armario, 2014). Dietary supplementation of SB leads to suppression of appetite through an action in the hypothalamus (Li et al., 2017). SB administration through drinking water promotes glucose metabolism in the brain at a dosage of 110 mg/kg/day (Val-Laillet et al., 2018). The mechanism of those actions is related to gene expression by the HDACi activity of SB. The gene expression-independent effect remains unknown (Stilling, van de Wouw, Clarke, Stanton, Dinan & Cryan, 2016). super-dosed SB generated a transient mitochondrial reprogramming in the brain for the reversible drop in body temperature. The effect was observed with mitochondrial swelling, crista loss, neurotransmitter reduction, and TCA metabolite accumulation in the brain hypothalamus. The brain inhibition was attenuated by NE, but not by CsA. CsA stabilizes the mitochondria in the cell culture, but not blocking the SB effects in the body for inability to cross the blood-brain barrier. These data suggest that the brain inhibition is the major cause of temperature drop by SB.

Current study revealed the pharmacological dynamics of SB in the brain tissue. SB reached the peak concentration of 58 μg/100g wet tissue in the brain, and stayed at the peak for 30 min with the half-life of 120 min. In contrast to the SB effects, the dynamics of SB concentration remains largely unknown in the brain (Stilling, van de Wouw, Clarke, Stanton, Dinan & Cryan, 2016). In a tissue distribution study, radiolabeled butyric acid (BA) was used in bamboo by vein injection. BA tissue concentrations were in following orders: pancreas, spleen, liver, kidney, vertebra and brain (Kim et al., 2013). With a half-life of 10 min in the blood, it takes days for pharmacological dose of SB to induce a stable effect through gene expression in the clinical application (Stilling, van de Wouw, Clarke, Stanton, Dinan & Cryan, 2016).

This study suggests that SB generated a metabolic reprogramming in the brain mitochondria independently of gene expression by HDACi. As a fuel supply, butyrate is consumed by mitochondria in production of ATP or heat (Donohoe et al., 2011; Li et al., 2019). Conversion of SB into acetyl-CoA is required for SB metabolism in mitochondria. However, the impact of super-dosed SB is unknown in the mitochondrial metabolism. The morphology change of mitochondria suggests that the mitochondrial membrane potential was lost in response to the SB challenge, which was observed with metabolite alteration in several pathways including TCA cycle and glycolysis, a result of mitochondrial metabolic reprogramming. The mechanism was investigated with the representative enzyme proteins in the respiratory chain and TCA cycles. No significant reduction was observed in those proteins although mRNA expression was changed for some of them. The reprogramming happened within 30 mins, which highly suggests a mechanism of post-translational modification or sterol regulation of the enzyme activities by SB. HDAC inhibitors including SAHA and sodium pyruvate failed to generate the same effect in the regulation of body temperature, suggesting that gene expression by HDACi may not play a major role in the preprogramming.

Current study demonstrates that the proton transportation activity of ANT proteins was induced by SB for the metabolic programming. The mitochondrial swelling and cristae loss was induced in the brain hypothalamic neurons. The structure change provides a mechanism for the mitochondrial paralysis and brain inhibition. Water influx into the mitochondrial matrix is a cause of the swelling, which is a result of mitochondrial permeability transient pore (mPTP) opening. Collapse of mitochondrial potential is a common trigger of the opening, which was observed in the SB-treated cells following proton leak. The SB-induced leak was inhibited by CsA and BKA (ANT inhibitor) in the cellular model, suggesting contribution of ANT activation to the mPTP opening. The possibility was proved in the experiment with shRNA-mediated gene knockdown of ANTs, which abolished the SB-induced proton leak. Both ANT isoforms (ANT1 and ANT2) were involved in the proton leak as the isoform-specific knockdown gave similar effects (Suppl. 4). These data suggest that ANT activation for proton transportation was induced by SB for the mPTP opening and mitochondrial structure change. The activation may a result of conformation change in the ANT proteins as the protein level was not elevated.

In summary, the super-dosed SB reduced the body energy metabolism with a reversible drop in the body temperature, which was attenuated by i.p. injection of norepinephrine. In the brain, the activity was associated with SB elevation and neurotransmitter reduction. SB triggered a rapid mitochondrial reprogramming in energy metabolism for the elevation in glycolysis, inhibition of TCA cycle and induction of proton leak. The thermogenesis of proton leak may warn up the heat-sensitive neurons in the hypothalamic circuit to trigger the temperature drop, which involves GABAergic neurons and TRPM2-expressing neurons (Song et al., 2016; Zhao et al., 2017). SB activated the proton transportation activity of ANT proteins for the proton leak without an increase in ANT proteins. The gene expression by histone deacetylase inhibition may not play a major role in the SB activity in the mitochondrial reprogramming.

## Author contributions

YX, SP, XC, SQ, SS, JL, HS, WS, XZ and WS conducted the experiments, analyzed the data, prepared the figures and manuscript draft. JY and WJ had the concept, designed the study, did data interpretation and manuscript writing. All authors have read and approved the manuscript.

## Acknowledgments

The project was supported by the National Key R&D Program of China (2018YFA0800603) and a project (19ZR1439000) of the Shanghai Association for Science and Technology to JY, and the internal fund of the Shanghai Sixth People’s Hospital to YX.

