## Supplementary material for "Super-dosed butyrate induces a revisable metabolic paralysis through transient mitochondrial reprogramming in the gut-brain axis": Suppllemental method

**Supplement 1**

**Methods**

**1. Mitochondrial membrane permeability**

After 1 hour of SB injection, the brain mitochondria were extracted as described above. The mitochondria potential was detected with JC-1 (2 μg/ml, 40705ES08, YEASEN, China) to evaluate the mitochondrial membrane permeability. The fluorescence signal of JC-1 was detected with a microplate reader (SpectraMaxi3x, Molecular Devices, USA) with excitation (550 nm) and emission (600 nm) wavelength. For the monomer, the excitation (485 nm) and the emission (535 nm) wavelength were used.

**2. qRT-PCR**

After 1 hour of SB injected, total RNA was extracted from the brain hemisphere using a TRIzol reagent, and mRNA was quantified following the reverse transcription into cDNA with a kit (HiScript® II Q RT SuperMix, R223, Vazyme Biotech, Nanjing, China) using 1 μg of RNA.18S was used as the reference. The primers were shown in the table below.

**3. Western Blot（WB）**

The proteins preparation, electrophoresis and membrane blotting were conducted as described elsewhere [1]. The protein sample (40 μg/sample) was loaded in each lane of the gel. The primary antibodies to CREB (#9197S), pCREB (Ser133，#9198S), HSL (#4107)，pHSL (Ser563，#4139), ANT2 (#14671), and acetylated-lysine (#9441) were purchased from the Cell Signaling Technology (Boston，USA). The primary antibodies to GAPDH (ab181602), ANT1 (ab110322), VDAC1 (ab154856), H3K27 (ab4729), TOMM20 (ab186735), Bad (ab32445), Bax (ab32503), Bcl-2 (ab182858), Bcl-xl (ab32370), Citrate synthase (ab129095), IDH2 (ab131263), OGDH (ab137773), SDHA (ab137040), MDH2 (ab181873), NDUFB8 (ab110242), Complex II (ab110410), UQCRC2 (ab14745), Complex IV (ab16056) were from the Abcam (Cambridge, England). Antibody to ATP5A1 (495240) was from Invitrogen (Camarillo, USA).

**4. Mitochondrial uncoupling assay in cultured cells**

The cell line Neuro-2a (CCL-131) was purchased from the American Type Culture Collection (ATCC, Manassas, VA, USA) , cultured in the complete medium of DMEM (SH30243.01, HyClone, Los Angeles, USA) supplemented with antibiotics (1%, GIBCO, USA) , 10% fetal bovine serum (FBS) (10270-106, GIBCO, Massachusetts, USA) and 0.5% GlutaMAX (35050-061, GIBCO, Massachusetts, USA) in the incubator of 37 ℃ and 5% CO_2_. For mitochondrial function, 2 x 10^4^ cells/well were seeded in the 24-well seahorse plate for 48 hours. The OCR was detected with the Seahorse XFe24. SB (20 mM) was administrated 1 hour before the test. The substrates contained 25 mM glucose, 1 mM pyruvate and 4 mM glutamine in the Base Medium (102353-100, Agilent, Santa Clara, USA). The activators and inhibitors were as used as mentioned above including Rotenone (0.5 μM). Uncoupling, proton leak and ATP production were calculated based on the changes in OCR.

| Name | Forward | Reverse |
| --- | --- | --- |
| 18S | 5-GCCGCTAGAGGTGAAATTCT-3 | 5-TCGGAACTACGACGGTATCT-3 |
| CS | 5-GCATGAAGGGACTTGTGTATGA-3 | 5-TTCTGGCACTCAGGGATACT-3 |
| IDH1 | 5-ATGCAAGGAGATGAAATGACACG-3 | 5-GCATCACGATTCTCTATGCCTAA-3 |
| IDH2 | 5-CCAGTACAAGGCCACAGATT-3 | 5-GTTATACACCTCCCACTCCTTG-3 |
| OGDH | 5-CACTTACCCACCACCACTTT-3 | 5-CCACTGGCATTGTTCCAAATC-3 |
| SDH | 5-GGCAAGTTCTGAGCCTGTA-3 | 5-CGGCGATACAGATACTCGATAC-3 |
| MDH2 | 5-GTTGCCAGATTGCCTCAAAG-3 | 5-CAATGGTAGCGTTGGTGTTG-3 |
| ANT1 | 5-CATCT ACAGAGCTGCCTACTTC-3 | 5-TCACACTCTGGGCAATCATC-3 |
| ANT2 | 5-TTGGTGACTGCCTGGTTAAG-3 | 5-GGCAGCTCGGTAGATAATG-3 |
| VDAC1 | 5-TGGACTGAAGCTCACCTTTG-3 | 5-GTTGATGTGCTCCCTCTTGT-3 |
| GPR41 | 5-CAGTGGCTGTGGACTTACTTT-3 | 5-GCAGAAGATGAAGGGCAGAA-3 |
| GPR43 | 5-CTACGAGAACTTCACCCAAGAG-3 | 5-GAAGCGCCAATAACAGAAGATG-3 |
| GPR109A | 5-CCTTATCTGGCTTCCACATCTC-3 | 5-GTTCAACGAACGGCCAAATC-3 |

Primer sequence for mouse genes examined by qRT-PCR
